# Design of the TRONCO BioConductor Package for TRanslational ONCOlogy

**DOI:** 10.1101/027524

**Authors:** Marco Antoniotti, Giulio Caravagna, Luca De Sano, Alex Graudenzi, Giancarlo Mauri, Bud Mishra, Daniele Ramazzotti

## Abstract

Models of *cancer progression* provide insights on the order of accumulation of genetic alterations during cancer development. Algorithms to infer such models from the currently available mutational profiles collected from different cancer patiens (*cross-sectional data*) have been defined in the literature since late 90s. These algorithms differ in the way they extract a *graphical model* of the events modelling the progression, e.g., somatic mutations or copy-number alterations.

TRONCO is an R package for TRanslational ONcology which provides a serie of functions to assist the user in the analysis of cross-sectional genomic data and, in particular, it implements algorithms that aim to model cancer progression by means of the notion of selective advantage. These algorithms are proved to outperform the current state-of-the-art in the inference of cancer progression models. TRONCO also provides functionalities to load input cross-sectional data, set up the execution of the algorithms, assess the statistical confidence in the results and visualize the models.

**Availability**. Freely available at http://www.bioconductor.org/ under GPL license; project hosted at http://bimib.disco.unimib.it/ and https://github.com/BIMIB-DISCo/TRONCO.

**Contact**.

## 1 Introduction

In the last two decades many specific genes and genetic mechanisms involved in different types of cancer have been identified. Yet our understanding of cancer and of its varied progressions is still largely elusive as it still faces fundamental challenges.

Meanwhile, a growing number of cancer-related genomic data sets have lately become available (e.g., see [9]). Thus, there now exists an urgent need to leverage a number of sophisticated computational methods in biomedical research to analyse such fast-growing biological datasets. Motivated by this state of affairs, we focus on the problem of *reconstructing progression models* of cancer. In particular, we aim at inferring the plausible sequences of *genomic alterations* that, by a process of *accumulation*, selectively make a tumor fitter to survive, expand and diffuse (i.e., metastasize).

We developed a number of algorithms (see [10, 12]) which are implemented in the *TRanslational ONCOlogy* (tronco) package. Starting from cross-sectional genomic data, such algorithms aim at reconstructing a probabilistic progression model by inferring “selectivity relations”, where a mutation in a gene *A* “selects” for a later mutation in a gene B. These relations are depicted in a combinatorial graph and resemble the way a mutation exploits its *“selective advantage*” to allow its host cells to expand clonally. Among other things, a selectivity relation implies a putatively invariant temporal structure among the genomic alterations (i.e., *events*) in a specific cancer type. In addition, a selectivity relation between a pair of events here signifies that the presence of the earlier genomic alteration (i.e., the *upstream event*) is advantageous in a Darwinian competition scenario raising the probability with which a subsequent advantageous genomic alteration (i.e., the *downstream event*) “survives” in the clonal evolution of the tumor (see [12]).

Notice that, in general, the inference of cancer progression models requires a complex data processing pipeline (see [3]), as summarized in Figure 1. Initially, one collects *experimental data* (which could be accessible through publicly available repositories such as TCGA) and performs *genomic analyses* to derive profiles of, e.g., somatic mutations or Copy-Number Variations for each patient. Then, statistical analysis and biological priors are used to select events relevant to the progression - e.g., *driver mutations*. This complex pipeline can also include further statistics and priors to determine cancer subtypes and to generate *patterns of selective advantage* -, e.g, hypotheses of mutual exclusivity. Given these inputs, our algorithms (such as CAPRESE and CAPRl) can extract a progression model and assess *confidence* measures using various metrics based on non-parametric bootstrap and hypergeometric testing. *Experimental validation* concludes the pipeline. TRONCO provides support to all the steps of the pipeline.

**Figure 1:**
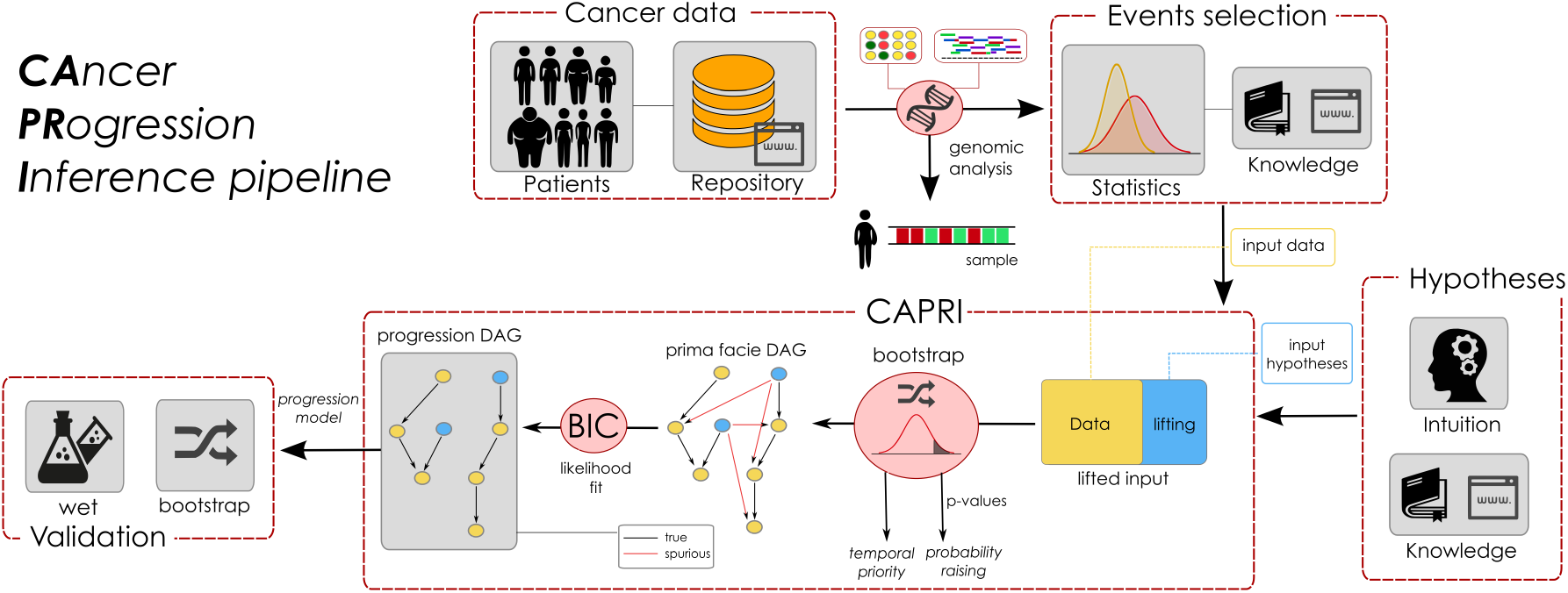
Data processing pipeline for the cancer progression inference. TRONCO implements a pipeline consisting in a series of functions and algorithms to extract cancer progression models from cross-sectional inout data. The first step of such a pipeline consists in collecting *experimental data* (which could be accessible through publicly available repositories such as TCGA) and performing *genomic analyses* to derive profiles of, e.g., somatic mutations or Copy-Number Variations for each patient or single cells. Then, both statistical analysis and biological priors are adopted to select the significant alterations for the progression; e.g., *driver mutations.* This complex pipeline can also include further statistics and priors to determine cancer subtypes and to generate *patterns of selective advantage*; e.g., hypotheses of mutual exclusivity. Given these inputs, the implemented algorithms (i.e., CAPRESE and capri) can extract a progression model and assess various *confidence* measures on its constituting relations such as non-parametric bootstrap and hypergeometric testing. *Experimental validation* concludes the pipeline, see [12] and [3].

### 2 Inference Algorithms

TRONCO, provides a series of functions to support the user in each step of the pipeline, i.e., from data import, through data visualization and, finally to the inference of cancer progression models. Specifically, in the current version, TRONCO implements CAPRESE and CAPRI algorithms for cancer progression inference, which we briefly describe in the following.

Central to these algorithms, is Suppes’ notion of *probabilistic causation*, which can be stated in the following terms: a selectivity relation between two observables *i* and *j* is said to hold if (1) *i* occurs earlier than *j* – *temporal priority* (TP) – and (2) if the probability of observing *i* raises the probability of observing *j*, i.e., 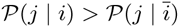 – *probability raising* (PR). For the detailed description of the methods, we refer the reader to [10, 12].

### 2.1 CAncer PRogression Extraction with Single Edge

The *CAncer PRogression Extraction with Single Edges* algorithm, i.e., caprese, extracts tree-based models of cancer progression with (*i*) multiple independent starting points and (*ii*) branches. The former models the emergence of different progressions as a result of the natural heterogeneity of cancer (cfr., [10]). The latter models the possibility of a clone to undergo positive selection by acquiring different mutations.

The inference of caprese’s models is driven by a *shrinkage* estimator of the confidence in the relation between pair of genes, which augments robustness to noise in the input data.

As shown in [10], caprese is currently the state-of-the-art algorithm to infer tree cancer progression models, although its expressivity is limited to this kind of selective advantage models (cfr., [12]). Since this limitation is rather unappealing in analyzing cancer data, an improved algorithm was sought in [12].

### 2.2 CAncer PRogression Inference

The *CAncer PRogression Inference* algorithm, i.e., capri, extends tree models by allowing multiple predecessors of any common downstream event, thus allowing construction of directed acyclic graph (DAGs) progression models.

capri performs maximum likelihood estimation for the progression model with constraints grounded in Supped’ prima facile causality (cfr., [12]). In particular, the search space of the possible valid solutions is limited to the selective advantage relations where both TP and PR are verified and then, on this reduced search space, the likelihood fit is performed.

In [12], capri was shown to be effective and polynomial in the size of the inputs.

### 2.3 Algorithms’ Structure

The core of the two algorithms is a simple quadratic loop^1^ that *prunes* arcs from an initially totally connected graph. Each pruning decision is based on the application of Suppes’ *probabilistic causation* criteria.

The pseudocode of the two implemented algorithms along with the procedure to evaluate the confidence of the arcs by bootstrap is summarized in Algorithms 1,2,3 and 4, which depict the data preparation step, the caprese and capri algorithms and finally the optional *bootstrap* step.

#### Algorithm 1: TRONCO Data Import and Preprocessing

**Input**: a data set containing `MAF` or `GISTIC` scores, e.g., as obtained from cBio portal ([4, 2]). Result: a data structure containing boolean flags for “events”, relative frequencies and other metadata.

1. From the dataset (depending on the data format) derive a Boolean matrix M, where each entry 〈i, *j*〉 is true if event *i* is “present” in sample/patient *j*.
2. **forall the** *events e* **do**
3. Compute the *frequency* of the event e in the dataset and save it in a map *F*.
4. Compute the *joint probability* of co-occurrence of pair of events in the dataset and save it in a map *C*.
5. **end**
6. **return** *A data structure comprising the Boolean matrix M, the maps F and C and other metadata.*

## 3 Package Design

In this section we will review the structure and implementation of the TRONCO package. For the sake of clarity, we will structure the description through the following functionalities that are implemented in the package.

- **Data import.** Functions for the importation of data both from flat files (e.g., MAF, GISTIC) and from Web querying (e.g., cBioPortal [4]).
- **Data export and correctness.** Functions for the export and visualization of the imported data.
- **Data editing.** Functions for the preprocessing of the data in order to tidy them.
- **External utilities.** Functions for the interaction with external tools for the analysis of cancer subtypes or groups of mutually exclusive genes.
- **Inference algorithms.** In the current version of TRONCO, the CAPRESE and CAPRI algorithms are provided in a polinomial implementation.
- **Confidence estimation.** Functions for the statistical estimation of the confidence of the reconstructed models.
- **Visualization.** Functions for the visualization of both the input data and the results of the inference and of the confidence estimation.

### Algorithm 2: CAPRESE algorithm

**Input:** a dataset of *n* events, i.e., genomic alterations, and *m* samples packed in a data structure obtained from Algorithm 1.

Result: a *tree model* representing all the relations of selective advantage.

*Pruning based on Suppes’ criteria.*

1. Let *G ←* a complete directed graph over the vertices *n*.
2. **forall the** *arcs* (*a, b*) *in G* **do**

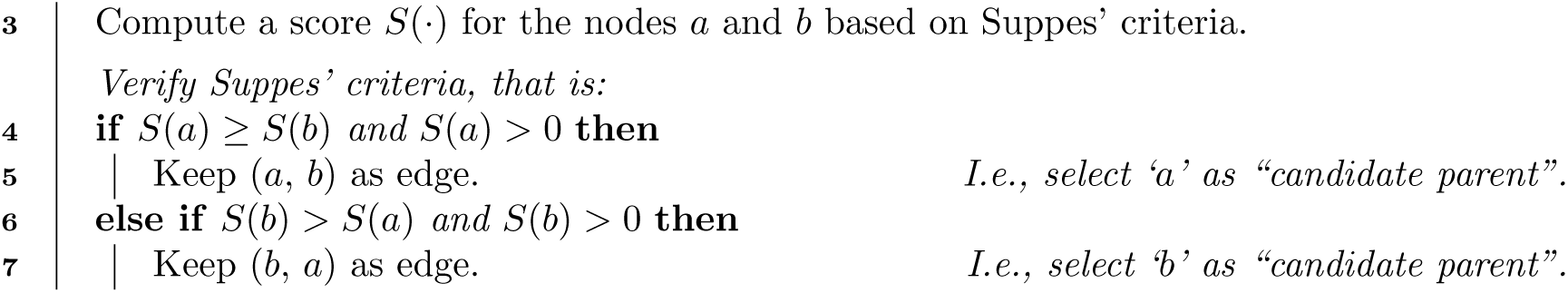

8 **end** *Fit of the* Prima Facie *directed acyclic graph to the best tree model.*
9 Let 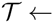 ← the best tree model obtained by Edmonds’ algorithm (see [5]).
10 Remove from 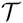 any connection where the candidate father does not have a minimum level of correlation with the child.
11 **return** *The resulting* tree *model* 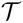.

### 3.1 Data Import

The starting point of tronco analysis pipeline, is a dataset of genomics alterations (i.e., somatic mutations and copy number variations) which need to be imported as a tronco compliant data structure, i.e., a R list structure containing the required data both for the inference and the visualization. The data import functions take as input such genomic data and from them create a tronco compliant data structure consisting in a list variable with the different parameters needed by the algorithms.

The core of data import from text files, is the function

~~~
import.genotypes(geno, event.type = “variant”, color = “Darkgreen”).
~~~

This function imports a matrix of 0/1 alterations as a TRONCO compliant dataset. The input geno can be either a dataframe or a file name. In any case the dataframe or the table stored in the file must have a column for each altered gene and a rows for each sample. Column names will be used to determine gene names; if data are loaded from a file, the first column will be assigned as row names.

tronco imports data from other file format such as MAF and GISTIC, by providing wrappers of the function `import.genotypes.` Specifically, the function

~~~
import.MAF(file, sep = “\t”, is.TCGA = TRUE)
~~~

imports mutation profiles from a *Manual Annotation Format* (MAF) file. All mutations are aggregated as a unique event type labeled `“Mutation”` and are assigned a color according to the default of function `import.genotypes.` If the input is in the TCGA MAF file format, the function also checks for multiple samples per patient and a warning is raised if any are found. The function

#### Algorithm 3: capri

**Input:** a dataset of *n* variables, i.e., genomic alterations or patterns, and *m* samples.

**Result:** a graphical model representing all the relations of “selective advantage”.

*Pruning based on the Suppes’ criteria*

1. Let *G* ← a directed graph over the vertices *n*
2. **forall the** *arcs (a, b)* ∈ *G* **do**
3. Compute a score *S*(.) for the nodes *a* and *b* in terms of Suppes’ criteria.
4. Remove the arc (*a*, *b*) if Suppes’ criteria are not met.
5. **end** *Likelihood fit on the Prima Facie directed acyclic graph*
6. Let 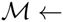 the subset of the remaining arcs *∈ G*, that maximize the log-likelihood of the model, computed as: 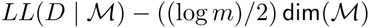 where *D* denotes the input data, *m* denotes the number of samples, and 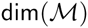 denotes the number of parameters in 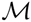 (see [8]).
7. **return** *The resulting* graphical *model* 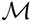.

~~~
import.GISTIC(x)
~~~

also transforms GISTIC scores for copy number alterations (CNAs) in a TRONCO compliant object. The input can be a matrix, with columns for each altered gene and rows for each sample; in this case colnames/rownames mut be provided. If the input is a character an attempt to load a table from file is performed. In this case the input table format should be consistent with TCGA data for focal CNA; i.e., there should hence be: one column for each sample, one row for each gene, a column `Hugo_Symbol` with every gene name and a column `Entrez_Gene_Id` with every genes Entrez ID. A valid GISTIC score should be any value of: `“Homozygous Loss” (−2), “Heterozygous Loss” (−1), “Low–level Gain” (+1), and “High-level Gain” (+2).`

Finally, tronco also provides utilities for the query of genomic data from cBioPortal [4]. This functionality is provided by the function

~~~
cbio.query(cbio.study = NA, cbio.dataset = NA, cbio.profile = NA, genes)
~~~

which is a wrapper for the `CGDS` package [7]. This can work either automatically, if one sets `cbio.study, cbio.dataset and cbio.profile,` or interactively. A list of genes to query with less than 900 entries should be provided. This function returns a list with two dataframes: the required genetic profile along with clinical data for the `cbio.study.` The output is also saved to disk as Rdata file. See also the cBioPortal page at http://www.cbioportal.org.

The function

~~~
show(x, view = 10)
~~~

prints (on the R console) a short report of a dataset x, which should be a TRONCO compliant dataset.

All the functions described in the following sections will assume as input a TRONCO compliant data structure.

### 3.2 Data Export and Correctness

tronco provides a series of function to explore the imported data and the inferred models. All these functions are named with the `“as.”` prefix.

#### Algorithm 4: Bootsrap Procedure

**Input**: a model 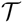 obtained from caprese or a model 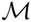 obtained from capri, and the initial dataset.

**Result**: the *confidence* in the inferred arcs.

1. Let *counter* ← 0
2. Let *nboot ←* the number of bootstrap sampling to be performed.
3. While *counter < nboot* **do**
4. Create a new dataset for the inference by random sampling of the input data.
5. Perform the reconstruction on the sampled dataset and save the results.
6. *counter = counter* + 1
7. **end**
8. Evaluate the confidence in the reconstruction by counting the number of times any arc is inferred in the sampled datasets.
9. **return** *The inferred model* 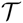 *or* 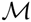 *augmented with an estimated confidence for each arc.*

Given a tronco compliant imported data set, the function

~~~
as.genotypes(x)
~~~

returns the 0/1 genotypes matrix. This function can be used in combination with the function

~~~
keysToNames(x, matrix)
~~~

to translate column names to event names, given the input matrix with colnames/rownames which represent genotypes keys. Also, functions to get the list of genes, events (i.e., each columns in the genotypes matrix, it differs from genes as the same genes of different types are considered different events), alterations (i.e., genes of different types are merged as 1 unique event), samples (i.e., patients or also single cells) and alteration types. See functions

~~~
as.genes(x, types = NA)
as.events(x, genes = NA, types = NA)
as.alterations(x, new.type = “Alteration”, new.color = “khaki”)
as.samples(x)
as.types(x, genes = NA).
~~~

Functions of this kind are also implemented to explore the results, most notably the models that have been inferred

~~~
as.models(x, models = names(x$model)))
~~~

the reconstructions

~~~
as.adj.matrix(x, events = as.events(x), models = names(x$model), type = “fit”)})
~~~

the patterns (i.e., the *formulæ*)

~~~
as.patterns(x)
~~~

and the confidence

~~~
as.confidence(x, conf).
~~~

Similarly, the library defines a set of functions that extract the cardinality of the compliant tronco data structure

~~~
nevents(x, genes = NA, types = NA)
ngenes(x, types = NA)
npatterns(x)
nsamples(x)
ntypes(x).
~~~

Furthermore, functions to asses the correctness of the inputs are also provided. The function

~~~
is.compliant(x,
               err.fun = “[ERR]”,
               stage = !(all(is.null(x$stages)) || all(is.na(x$stages))))
~~~

verifies that the parameter x is a compliant data structure. The function

~~~
consolidate.data(x, print = FALSE)
~~~

verifies if the input data are consolidated, i.e., if there are events with 0 or 1 probability or indistinguishable in terms of observations. Any indistinguishable event is returned by the function `duplicates(x).`

Finally, tronoo provides functions to access TCGA data.

~~~
TCGA.multiple.samples(x)
~~~

checks if whether are multiple sample in the input, while

~~~
TCGA.remove.multiple.samples(x)
~~~

removes them accordingly to TCGA barcodes naming rules.

### 3.3 Data Editing

tronoo provides a wide range of editing functions. We will describe some of them in the following; for a technical description we refer to the manual.

#### 3.3.1 Removing and Merging

A set of functions to remove items from the data is provided; such functions are characterized by the `delete.` prefix. The main functions are

~~~
delete.gene(x, gene)
delete.samples(x, samples)
delete.type(x, type)
delete.pattern(x, type)
~~~

that respectively remove genes, samples (i.e., tumors profiles), types (i.e., type of alteration such as somatic mutation, copy number alteratio, etc.), and patterns from a tronoo data structure x. Conversely it is possible to *merge* events and types:

~~~
merge.events(x, …, new.event, new.type, event.color)
merge.types(x, …, new.type = “new.type”, new.color = “khaki”)}).
~~~

#### 3.3.2 Binding

The purpose of the binding functions is to combine different datasets. The function

ebind(…)

combines events from one or more datasets, whose events need be defined over the same set of samples. The function

sbind(…)

combines samples from one or more datasets, whose samples need to be defined over the same set of events. Samples and events of two dataset can also be intersected via the function

~~~
intersect.datasets(x, y, intersect.genomes = TRUE).
~~~

#### 3.3.3 Changing and Renaming

The functions

~~~
rename.gene(x, old.name, new.name)
rename.type(x, old.name, new.name)
~~~

can be used respectively to rename genes or alterations types.

The function

~~~
change.color(x, type, new.color)
~~~

can be used to change the color associated to the specified alteration type in x.

#### 3.3.4 Selecting and Splitting

Genomics data usually involve a large number of genes, most of which are not relevant for cancer development (e.g., the may be passenger mutations). For this reason, tronco implements the function

~~~
events.selection(x, filter.freq = NA, filter.in.names = NA,filter.out.names = NA)
~~~

which allows the user to select a subset of genes to be analyzed. The selection can be performed by frequency and gene symbols. The 0 probability events can are removed by the function `trim(x).` Moreover, the functions

~~~
samples.selection(x, samples)
ssplit(x, clusters, idx = NA)
~~~

respectively filter a dataset x based on selected samples id and split the dataset into clusters (i.e., groups). The last function can be used to analyze specific subtypes within a tumor.

### 3.4 External Utilities

tronco permits the interaction with external tools to (*i*) reduce inter-tumor heterogeneity by cohort subtyping and (*ii*) detect fitness equivalent exclusive alterations. The first issue can be attacked by adopting clustering techniques to split the dataset in order to analyze each cluster subtype separately. Currently, TRONCO can export and inport data from [6] via the function

~~~
export.nbs.input(x, map\_hugo\_entrez, file = “tronco\_to\_nbs.mat”)
~~~

and the previously described splitting functions.

In order to handle alterations with equivalent fitness, TRONCO interacts with the tool MUTEX proposed in [1]. The interaction is ensured by the functions

~~~
export.mutex(x,
             filename = “to_mutex”,
             filepath = “./”,
             label.mutation = “SNV”,
             label.amplification = list(“High-level Gain”),
             label.deletion = list(“Homozygous Loss”))
import.mutex.groups(file, fdr = 0.2, display = TRUE)
~~~

. Such exclusivity groups can then be further added as patterns (see the next section).

### 3.5 Inference Algorithms

The current version of TRONCO implements the *progression reconstruction algorithms* algorithms caprese [10] and capri [12].

**caprese**. The caprese algorithm [10] can be executed by the function

~~~
tronco.caprese(data, lambda = 0.5, do.estimation = FALSE, silent = FALSE)
~~~

with data being a tronco data structure. The parameter `lambda` can be used to tune the shrinkage-like estimator adopted by CAPRESE, with the default being 0.5 as suggested in [10].

**capri**. The CAPRI algorithm [12] is executed by the function

~~~
tronco.capri(data,
                command = “hc”,
                regularization = c(“bic”, “aic”),
                do.boot = TRUE,
                nboot = 100,
                pvalue = 0.05,
                min.boot = 3,
                min.stat = TRUE,
                boot.seed = NULL,
                do.estimation = FALSE,
                silent = FALSE)
~~~

with data being a tronco data structure. The parameters `command` and `regularization` allow respectively to choose the heuristic search to be performed to fit the network and the regularizer to be used in the likelihood fit (see [12]). CAPRI can be also executed with or without the bootstrap preprocessing step depending on the value of the parameter `do.boot;` this is discouraged, but can speed up the execution with large input datasets.

As discussed in [12], capri constrains the search space using Suppes’ prima facie conditions which lead to a subset of possible valid selective advantage relations. The members of this subset are then evaluated by the likelihood fit. Although uncommon, it may so happen (especially when patterns are given as input) that such a resulting prima facie graphical structure may still contain cycles. When this happens, the cycles are removed through the heuristic algorithm implemented in

~~~
remove.cycles(adj.matrix,
             weights.temporal.priority,
             weights.matrix,
             not.ordered,
             hypotheses = NA,
             silent).
~~~

The function takes as input a set of weights in term of confidence for any selective advantage valid edge, ranks all the valid edges in increasing confidence levels and, starting from the less confident, goes through each edge removing the ones that can break the cycles.

#### 3.5.1 Patterns

capri allows for the input of patterns, i.e., group of events which express possible selective advantage relations. Such patterns are given as input using the function

~~~
hypothesis.add(data,
               pattern.label,
               lifted.pattern,
               pattern.effect = “*”,
               pattern.cause = “*”).
~~~

This function is wrapped within the functions

~~~
hypothesis.add.homologous(x,
                          pattern.cause = “*”,
                          pattern.effect = “*”,
                          genes = as.genes(x),
                          FUN = OR)
~~~

~~~
hypothesis.add.group(x,
                         FUN,
                         group,
                         pattern.cause = “*”,
                         pattern.effect = “*”,
                         dim.min = 2,
                         dim.max = length(group),
                         min.prob = 0)
~~~

which, respectively, allow the addition of analogous patterns (i.e., patterns involving the same gene of different types) and patterns involving a specified group of genes. In the current version of tronco, the implemented patterns are Boolean, i.e., those expressible by the boolean operators AND, OR and XOR (functions `AND(…), OR(…)` and `XOR(…)).`

### 3.6 Confidence Estimation

To asses the confidence of the selectivity relations found, tronco uses *non-parametric* and *statistical* bootstraps. For the non-parametric bootstrap, each event row is uniformly sampled with repetitions from the input genotype and then, on such an input, the inference algorithms are performed. The assessment concludes after *K* repetitions (e.g., *K* = 100). Similarly, for capri, a statistical bootstrap is provided: in this case the input dataset is kept fixed, but different seeds for the statistical procedures are sampled (see, e.g., [13] for an overview of these methods). The bootstrap is implemented in the function

~~~
tronco.bootstrap(reconstruction,
                 type = “non-parametric”,
                 nboot = 100,
                 verbose = FALSE)
~~~

where `reconstruction` is a tronco compliant object obtained by the inference by one of the implemented algorithms.

### 3.7 Visualization and Reporting

During the development of the tronco package, a lot of attention was paid to the visualization features which are crucial for the understanding of biological results. Listed below is a summary of the main features; for a detailed description of each function, please refer to the manual.

#### OncoPrint

OncoPrints are compact means of visualizing distinct genomic alterations, including somatic mutations, copy number alterations, and mRNA expression changes across a set of cases. They are extremely useful for visualizing gene set and pathway alterations across a set of cases, and for visually identifying trends, such as trends in mutual exclusivity or cooccurence between gene pairs within a gene set. Individual genes are represented as rows, and individual cases or patients are represented as columns. See http://www.cbioportal.org/. The function

~~~
oncoprint(x)
~~~

provides such visualizations with a TRONCO compliant data structure as input. The function `oncoprint.cbio(x)`

exports the input for the cBioPortal visualization, see http://www.cbioportal.org/public-portal/oncoprinter.jsp.

It is also possible to annotate a description and tumor stages to any oncoprint by means of the functions

~~~
annotate.description(x, label)
annotate.stages(x, stages, match.TCGA.patients = FALSE).
~~~

#### Reconstruction

The inferred models can be displayed by the function `tronco.plot.` The features included in the plots are multiple, such as the choice of the regularizer(s), editing font of nodes and edges, scaling nodes’ size in terms of estimated marginal probabilities, annotating the pathway of each gene and displaying the estimated confidence of each edge. We refer to the manual for a detailed description.

#### Reports

Finally, tronco provides a number of reporting utilities. The function

~~~
genes.table.report(x,
                   name,
                   dir = getwd(),
                   maxrow = 33,
                   font = 10,
                   height = 11,
                   width = 8.5,
                   fill = “lightblue”)
~~~

can be used to generate 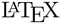 code to be used as report, while the function

~~~
genes.table.plot(x, name, dir = getwd())
~~~

generates histograms reports.

## 4 tronco Use Cases

In this Section, we will present a case study for the usage of the tronco package based on the work presented in [12]. Specifically, the example is from [11] where Piazza *et al.* used high-throughput *exome sequencing technology* to identity somatically acquired mutations in 64 aCML patients, and found a previously unidentified recurring *missense point mutation* hitting Setbp1.

The example illustrates the typical steps that are necessary to perform a *progression reconstruction* with tronco. The steps are the following:

1. Selecting “Events”.
2. Adding “Hypotheses”.
3. Reconstructing the “Progression Model”.
4. Bootstrapping the Data.

(In the following, user input at the console is shown in `boldface.`

### Selecting Events

We will start by loading the tronco package in R along with an example *dataset* that is part of the package distribution.

~~~
> library(TRONCO)
> data(aCML)
> hide.progress.bar <<- TRUE
~~~

We then use the function `show` to get a short summary of the aCML dataset that has just been loaded.

~~~
> show(aCML)
Description: CAPRI – Bionformatics aCML data.
Dataset: n=64, m=31, |G|=23.
Events (types): Ins/Del, Missense point, Nonsense Ins/Del, Nonsense point.
Colors (plot): darkgoldenrod1, forestgreen, cornflowerblue, coral.
Events (10 shown):
  gene 4: Ins/Del TET2
  gene 5: Ins/Del EZH2
  gene 6: Ins/Del CBL
  gene 7: Ins/Del ASXL1
  gene 29: Missense point SETBP1
  gene 30: Missense point NRAS
  gene 31: Missense point KRAS
  gene 32: Missense point TET2
  gene 33: Missense point EZH2
  gene 34: Missense point CBL
Genotypes (10 shown):
~~~

~~~
           gene 4      gene 5    gene 6     gene 7    gene 29    gene 30    gene 31    gene 32    gene 33    gene 34
patient 1      0      0      0      0      1      0      0      0      0      0
patient 2      0      0      0      0      1      0      0      0      0      1
patient 3      0      0      0      0      1      1      0      0      0      0
patient 4      0      0      0      0      1      0      0      0      0      1
patient 5      0      0      0      0      1      0      0      0      0      0
patient 6      0      0      0      0      1      0      0      0      0      0
~~~

Using the function `as.events,` we can have a look at the genes flagged as “mutated” in the dataset (i.e., the *events* that tronco deals with).

~~~
> as.events(aCML)
~~~

~~~
             type event
gene 4       "Ins/Del"               "TET2"
gene 5       "Ins/Del"                EZH2"
gene 6       "Ins/Del"               "CBL"
gene 7       "Ins/Del"               "ASXL1"
gene 29     "Missense point"       "SETBP1"
gene 30     "Missense point"       "NRAS"
gene 31     "Missense point"       "KRAS"
gene 32     "Missense point"     "TET2"
gene 33     "Missense point"     "EZH2"
…
gene 88     "Nonsense     point"     "TET2"
gene 89     "Nonsense     point"     "EZH2"
gene 91     "Nonsense     point"     "ASXL1"
gene 111     "Nonsense     point"     "CSF3R"
~~~

These events account for alterations in the following genes.

~~~
> as.genes(aCML)
[1]     "TET2"     "EZH2"     "CBL"     "ASXL1"     "SETBP1"     "NRAS"     "KRAS"     "IDH2"     "SUZ12"
[10]     "SF3B1"     "JARID2"     "EED"     "DNMT3A"     "CEBPA"     "EPHB3"     "ETNK1"     "GATA2"     "IRAK4"
[19]     "MTA2"     "CSF3R"     "KIT"     "WT1"     "RUNX1"
~~~

Now we can take a look at the alterations of only the gene SETBP1 across the samples.

~~~
> as.gene(aCML, genes = 'SETBP1')
     Missense point SETBP1
patient 1          1
patient 2          1
patient 3          1
…
patient 12          1
patient 13          1
patient 14          1
patient 15          0
patient 16          0
patient 17          0
…
patient 62          0
patient 63          0
patient 64          0
~~~

We consider a subset of all the genes in the dataset to be involved in patterns based on the support we found in the literature. See [12] as a reference.

~~~
> gene.hypotheses = c(‘KRAS’, ‘NRAS’, ‘IDH1’, ‘IDH2’, ‘TET2’, ‘SF3B1’, ‘ASXL1’)
~~~

Regardless from which types of mutations we include, we select only the genes which appear alterated in at least 5% of the patients. Thus, we first transform the dataset into *“alterations*” (i.e., collapsing all the event types for the same gene), and then we consider only these events from the original dataset.

~~~
> alterations = events.selection(as.alterations(aCML), filter.freq = .05)
*** Aggregating events of type(s) Ins/Del, Missense point, Nonsense Ins/Del, Nonsense point in a unique event with label “Alteration”.
Dropping event types Ins/Del, Missense point, Nonsense Ins/Del, Nonsense point for 23 genes.
*** Binding events for 2 datasets.
*** Events selection: #events=23, #types=1 Filters freq|in|out = {TRUE, FALSE, FALSE}
Minimum event frequency: 0.05 (3 alterations out of 64 samples).
Selected 7 events.
~~~

Selected 7 events, returning.

We now show a plot of the selected genes. Note that this plot has no title as by default the function events.selection does not add any. The resulting figure is shown in 2.

**Figure 2:**
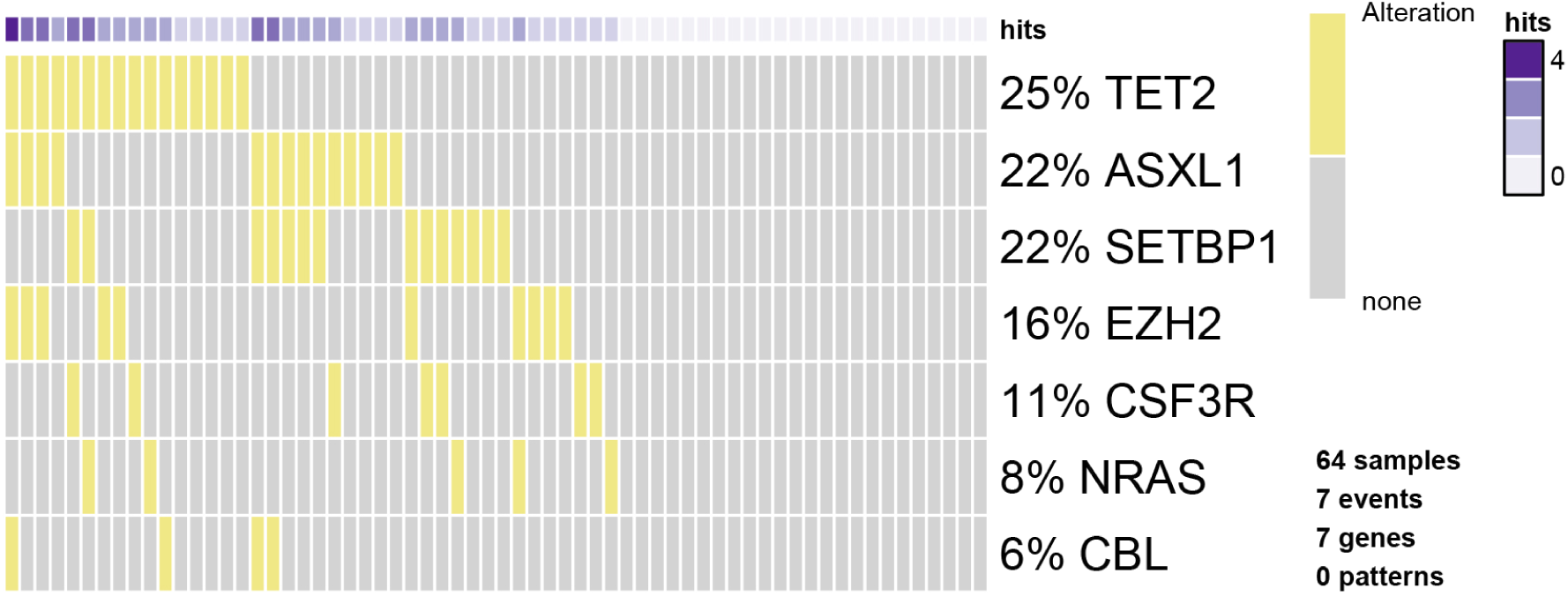
Oncoprint function in tronco. Result of the oncoprint function in tronco on the aCML dataset.

**Figure 3:**
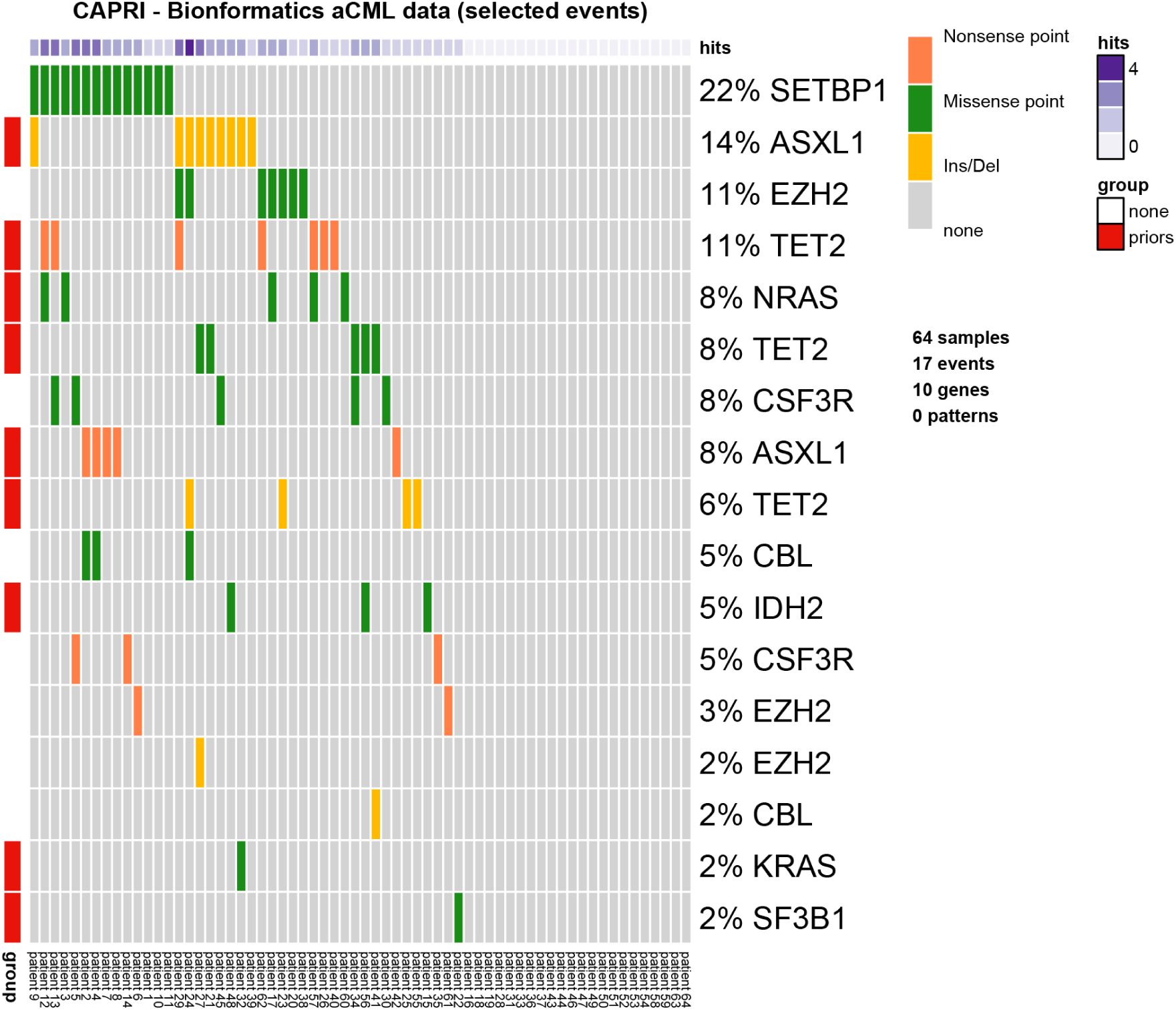
Annotated oncoprint. Result of the oncoprint function on the selected dataset in tronco with annotations.

~~~
> oncoprint(alterations,font.row = 12, cellheight = 20, cellwidth = 4)
*** Oncoprint for “”
with attributes: stage=FALSE, hits=TRUE
Sorting samples ordering to enhance exclusivity patterns.
~~~

### Adding Hypotheses

We now create the *dataset* to be used for the inference of the progression model. We consider the original dataset and from it we select all the genes whose mutations are occurring at least 5% of the times together with any gene involved in any hypothesis. To do so, we use the parameter `filter.in.names` as shown below.

~~~
> hypo = events.selection(aCML,
                            filter.in.names = c(as.genes(alterations),
                            gene.hypotheses))
*** Events selection: #events=31, #types=4 Filters freq|in|out = {FALSE, TRUE, FALSE} [filter.in] Genes hold: TET2, EZH2, CBL, ASXL1, SETBP1 … [10/14 found].
Selected 17 events, returning.
> hypo = annotate.description(hypo, ‘CAPRI – Bionformatics aCML data (selected events)’)
~~~

We show a new oncoprint of this latest dataset where we annotate the genes in gene.hypotheses in order to identify them 3. The sample names are also shown.

~~~
> oncoprint(hypo,
                  gene.annot = list(priors = gene.hypotheses),
                  sample.id = T,
                  font.row = 12,
                  font.column = 5,
                  cellheight = 20,
                  cellwidth = 4)
*** Oncoprint for “CAPRI – Bionformatics aCML data (selected events)”
with attributes: stage=FALSE, hits=TRUE
Sorting samples ordering to enhance exclusivity patterns.
Annotating genes with RColorBrewer color palette Set1.
~~~

We now also add the hypotheses that are described in CAPRI’s manuscript. Hypothesis of hard exclusivity (XOR) for NRAS/KRAS events (Mutation). This hypothesis is tested against all the events in the dataset.

~~~
> hypo = hypothesis.add(hypo, ‘NRAS xor KRAS’, XOR(‘NRAS’, ‘KRAS’))
~~~

We then try to include also a soft exclusivity (OR) pattern but, since its “*signature*” is the same of the hard one just included, it will not be included. The code below is expected to result in an error.

~~~
> hypo = hypothesis.add(hypo, ‘NRAS or KRAS’, OR(‘NRAS’, ‘KRAS’))
Error in hypothesis.add(hypo, "NRAS or KRAS", OR("NRAS", "KRAS")):
[ERR] Pattern duplicates Pattern NRAS xor KRAS.
~~~

To better highlight the perfect (hard) exclusivity among NRAS/KRAS mutations, one can examine further their alterations. See Figure 4.

**Figure 4:**
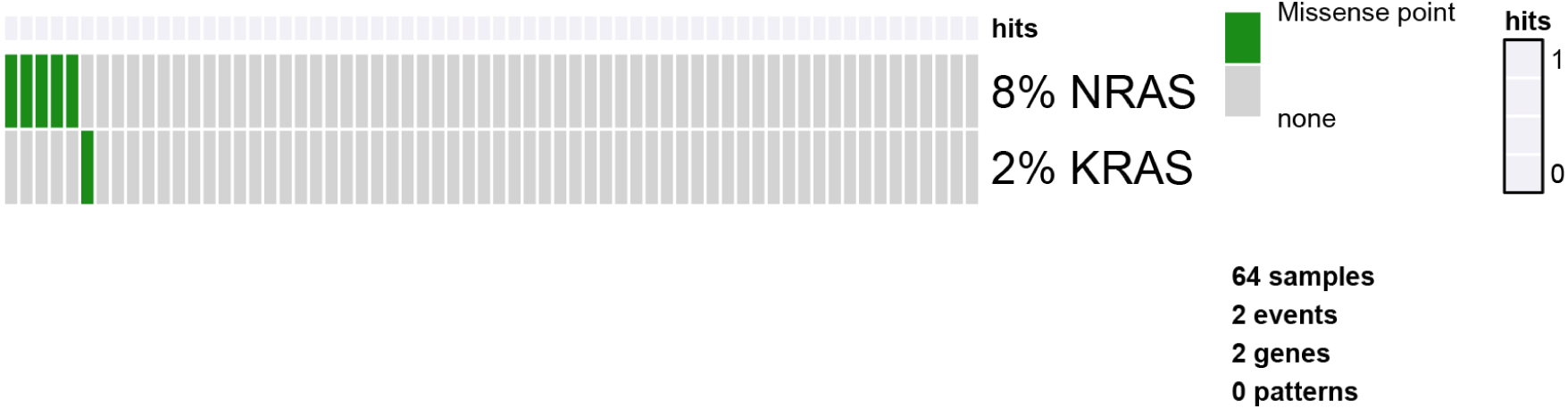
RAS oncoprint. Result of the oncoprint function in tronco for only the RAS genes to better show their hard exclusivity pattern.

~~~
> oncoprint(events.selection(hypo,
                           filter.in.names = c(‘KRAS’, ‘NRAS’)),
                           font.row = 12,
                           cellheight = 20,
                           cellwidth = 4)
*** Events selection: #events=18, #types=4 Filters freq|in|out = {FALSE, TRUE, FALSE}
[filter.in] Genes hold: KRAS, NRAS … [2/2 found].
Selected 2 events, returning.
*** Oncoprint for “”
with attributes: stage=FALSE, hits=TRUE
Sorting samples ordering to enhance exclusivity patterns.
~~~

We repeated the same analysis as before for other hypotheses and for the same reasons, we will include only the hard exclusivity pattern.

~~~
> hypo = hypothesis.add(hypo, ‘SF3B1 xor ASXL1’, XOR(‘SF3B1’, OR(‘ASXL1’)), ‘*’)
> hypo = hypothesis.add(hypo, ‘SF3B1 or ASXL1’, OR(‘SF3B1’, OR(‘ASXL1’)), ‘*’)
Error in hypothesis.add(hypo, “SF3B1 or ASXL1”, OR(“SF3B1”, OR(“ASXL1”)),:
[ERR] Pattern duplicates Pattern SF3B1 xor ASXL1.
~~~

Finally, we now repeat the same for genes TET2 and IDH2. In this case 3 events for the gene TET2 are present: “Ins/Del”, “Missense point” and “Nonsense point”. For this reason, since we are not specifying any subset of such events to be considered, all TET2 alterations are used. Since the events present a perfect hard exclusivity, their patterns will be included as a XOR. See Figure 5.

**Figure 5:**
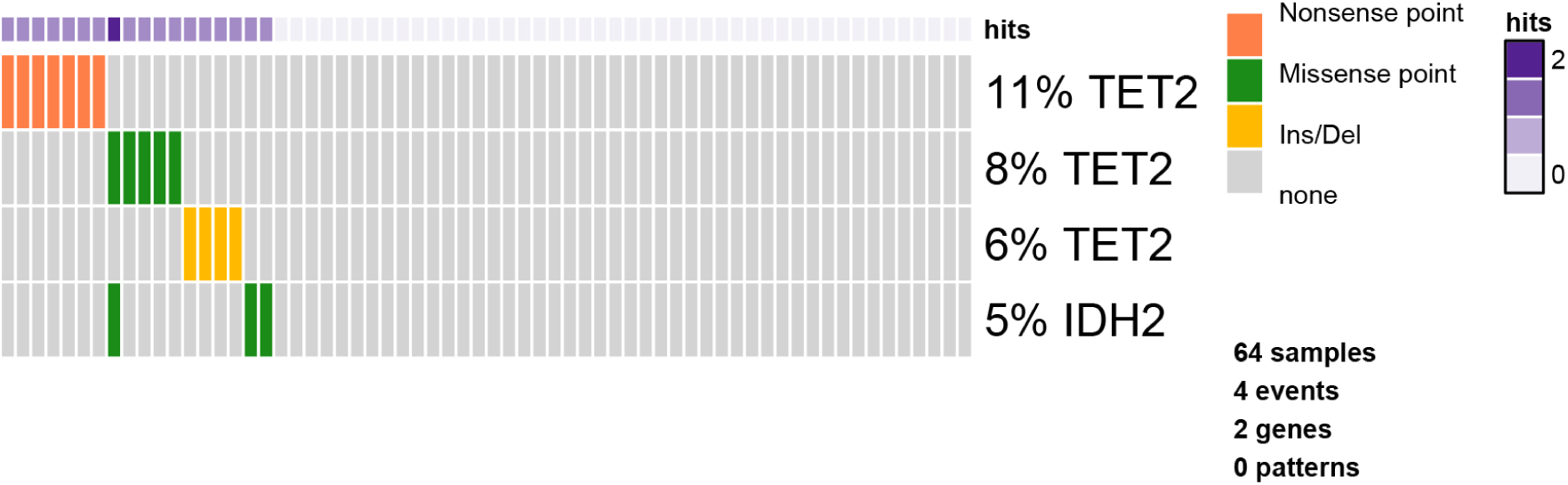
TET/IDH2 oncoprint. Result of the oncoprint function in tronco for only the TET/IDH2 genes.

~~~
> as.events(hypo, genes = ‘TET2’)
type event
gene 4 “Ins/Del”                 “TET2”
gene 32 “Missense point”   “TET2”
gene 88 “Nonsense point”   “TET2”
> hypo = hypothesis.add(hypo, ‘TET2 xor IDH2’, XOR(‘TET2’, ‘IDH2’), ‘*’)
> hypo = hypothesis.add(hypo, ‘TET2 or IDH2’, OR(‘TET2’, ‘IDH2’), ‘*’)
> oncoprint(events.selection(hypo, filter.in.names = c(‘TET2’, ‘IDH2’)),font.row=12, cellheight=20,cellwidth=4)
*** Events selection: #events=21, #types=4 Filters freq|in|out = {FALSE, TRUE, FALSE} [filter.in] Genes hold: TET2, IDH2 … [2/2 found].
Selected 4 events, returning.
*** Oncoprint for “”
with attributes: stage=FALSE, hits=TRUE
Sorting samples ordering to enhance exclusivity patterns.
~~~

We now finally add any possible group of homologous events. For any gene having more than one event associated we also add a soft exclusivity pattern among them.

~~~
> hypo = hypothesis.add.homologous(hypo)
*** Adding hypotheses for Homologous Patterns
Genes: TET2, EZH2, CBL, ASXL1, CSF3R
Function: OR
Cause: *
Effect: *
Hypothesis created for all possible gene patterns.
~~~

The final dataset that will be given as input to CAPRI is now finally shown. See Figure 6.

**Figure 6:**
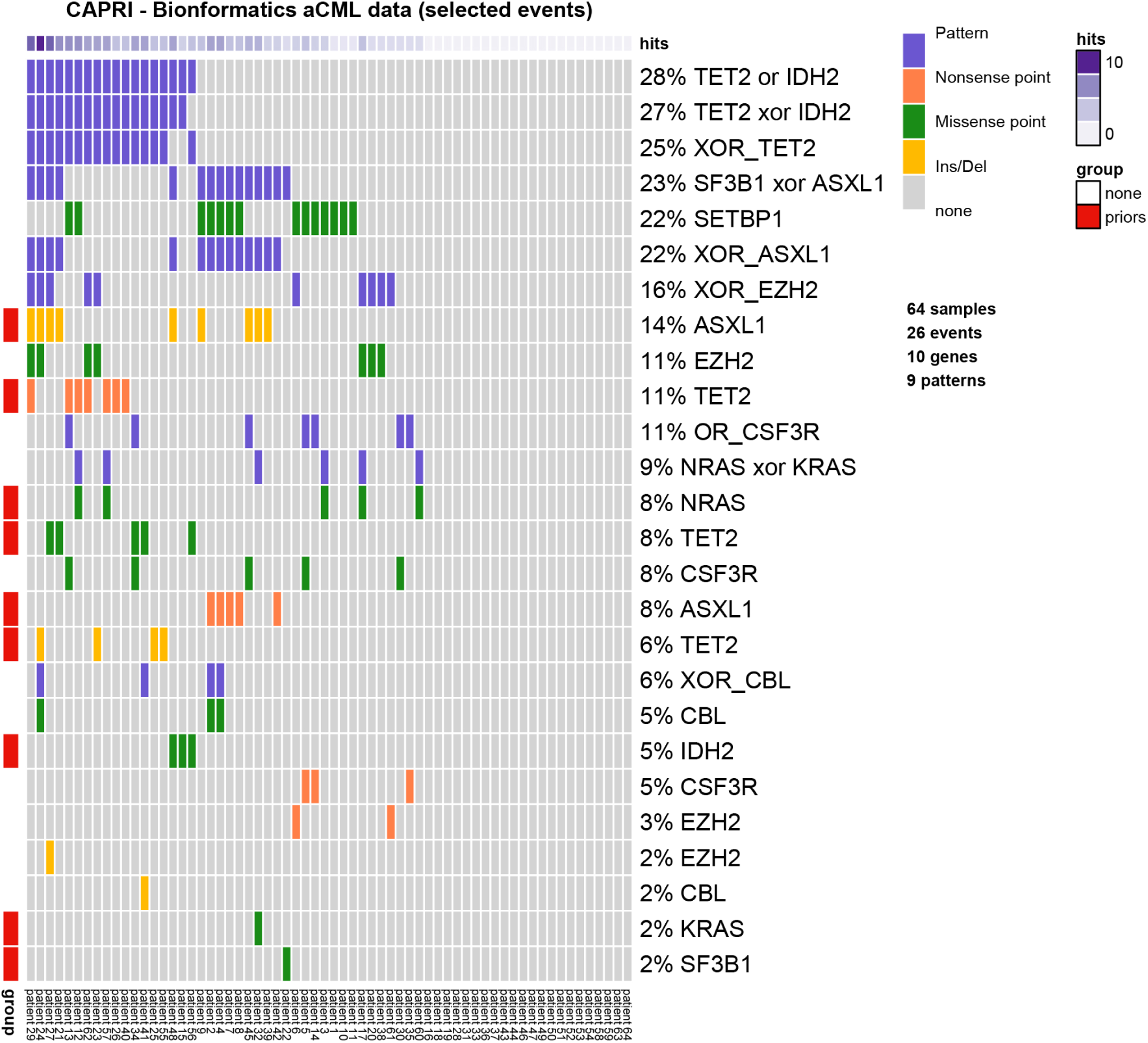
Final dataset for capri. Result of the oncoprint function in tronco on the dataset used in [12].

~~~
> oncoprint(hypo,
              gene.annot = list(priors = gene.hypotheses),
              sample.id = T,
              font.row = 10,
              font.column = 5,
              cellheight = 15,
              cellwidth = 4)
*** Oncoprint for “CAPRI – Bionformatics aCML data (selected events)”
with attributes: stage=FALSE, hits=TRUE
Sorting samples ordering to enhance exclusivity patterns.
Annotating genes with RColorBrewer color palette Set1.
~~~

### Reconstructing Progression Models

We next infer the model by running CAPRI algorithm with its default parameters: we use both AIC and BIC as regularizers, Hill-climbing as heuristic search of the solutions and exhaustive bootstrap (nboot replicates or more for Wilcoxon testing, i.e., more iterations can be performed if samples are rejected), p-value set at 0.05. We set the seed for the sake of reproducibility.

~~~
> model = tronco.capri(hypo, boot.seed = 12345, nboot = 10)
*** Checking input events.
*** Inferring a progression model with the following settings.
Dataset size: n = 64, m = 26.
Algorithm: CAPRI with “bic, aic” regularization and “hc” likelihood-fit strategy.
Random seed: 12345.
Bootstrap iterations (Wilcoxon): 10.
exhaustive bootstrap: TRUE.
p-value: 0.05.
minimum bootstrapped scores: 3.
*** Bootstraping selective advantage scores (prima facie).
Evaluating “temporal priority” (Wilcoxon, p-value 0.05)
Evaluating “probability raising” (Wilcoxon, p-value 0.05)
*** Loop detection found loops to break.
Removed 26 edges out of 68 (38%)
*** Performing likelihood-fit with regularization bic.
*** Performing likelihood-fit with regularization aic.
The reconstruction has been successfully completed in 00h:00m:02s
~~~

We then plot the model inferred by capri with BIC as a regularizer and we set some parameters to get a good plot; the confidence of each edge is shown both in terms of temporal priority and probability raising (selective advantage scores) and hypergeometric testing (statistical relevance of the dataset of input). See Figure 7.

**Figure 7:**
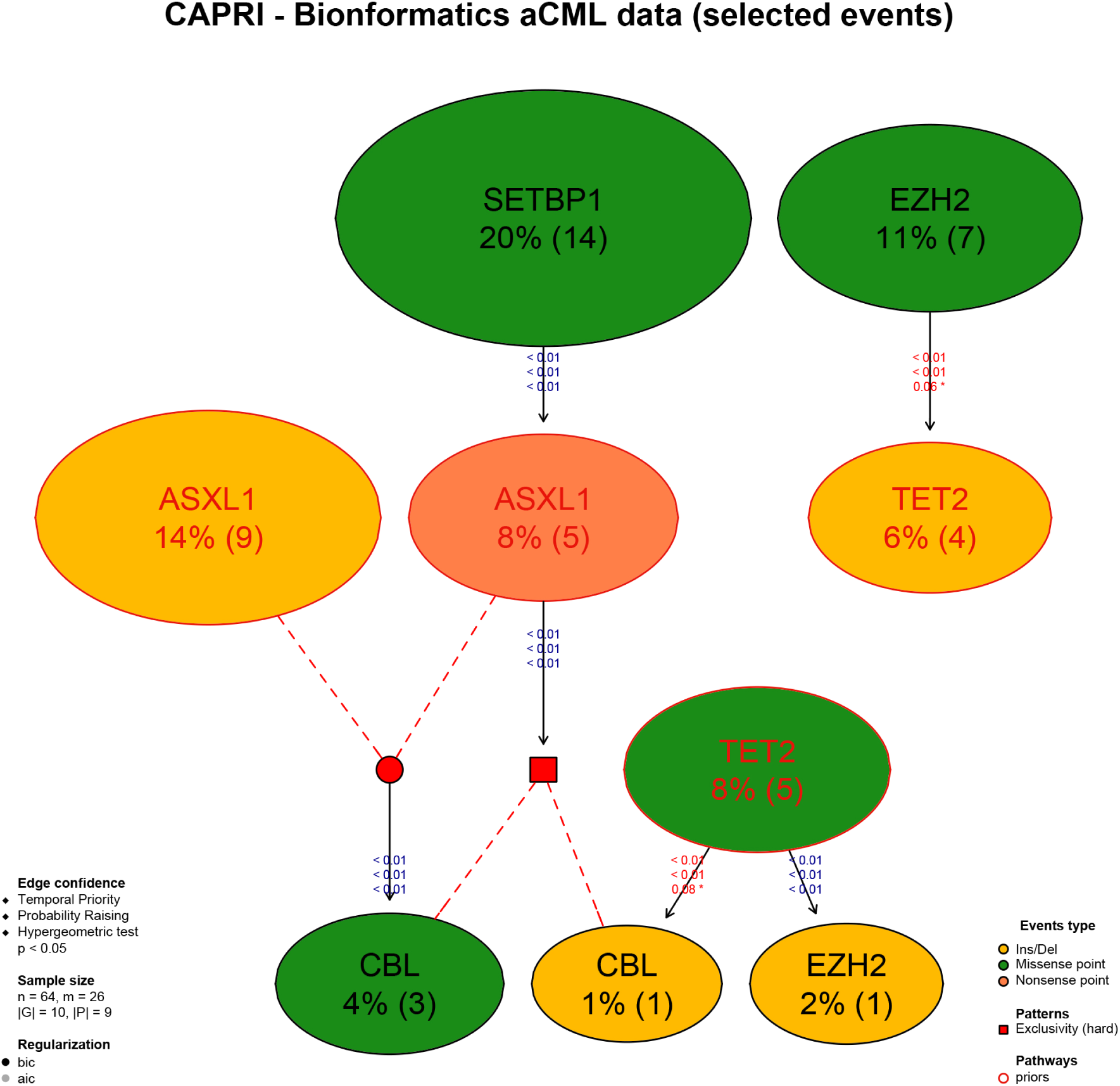
Reconstruction by capri. Result of the reconstruction by CAPRI on the input dataset.

~~~
> tronco.plot(model,
                 fontsize = 13,
                 scale.nodes = .6,
                 regularization = “bic”,
                 confidence = c(‘tp’, ‘pr’, ‘hg’),
                 height.logic = 0.25,
                 legend.cex = .5,
                 pathways = list(priors = gene.hypotheses),
                 label.edge.size = 5)
*** Expanding hypotheses syntax as graph nodes:
*** Rendering graphics
Nodes with no incoming/outgoing edges will not be displayed.
Annotating nodes with pathway information.
Annotating pathways with RColorBrewer color palette Set1.
Adding confidence information: tp, pr, hg
RGraphviz object prepared.
Plotting graph and adding legends.
~~~

### Bootstrapping the Data

Finally, we perform non-parametric bootstrap as a further estimation of the confidence in the inferred results. See Figure 8.

**Figure 8:**
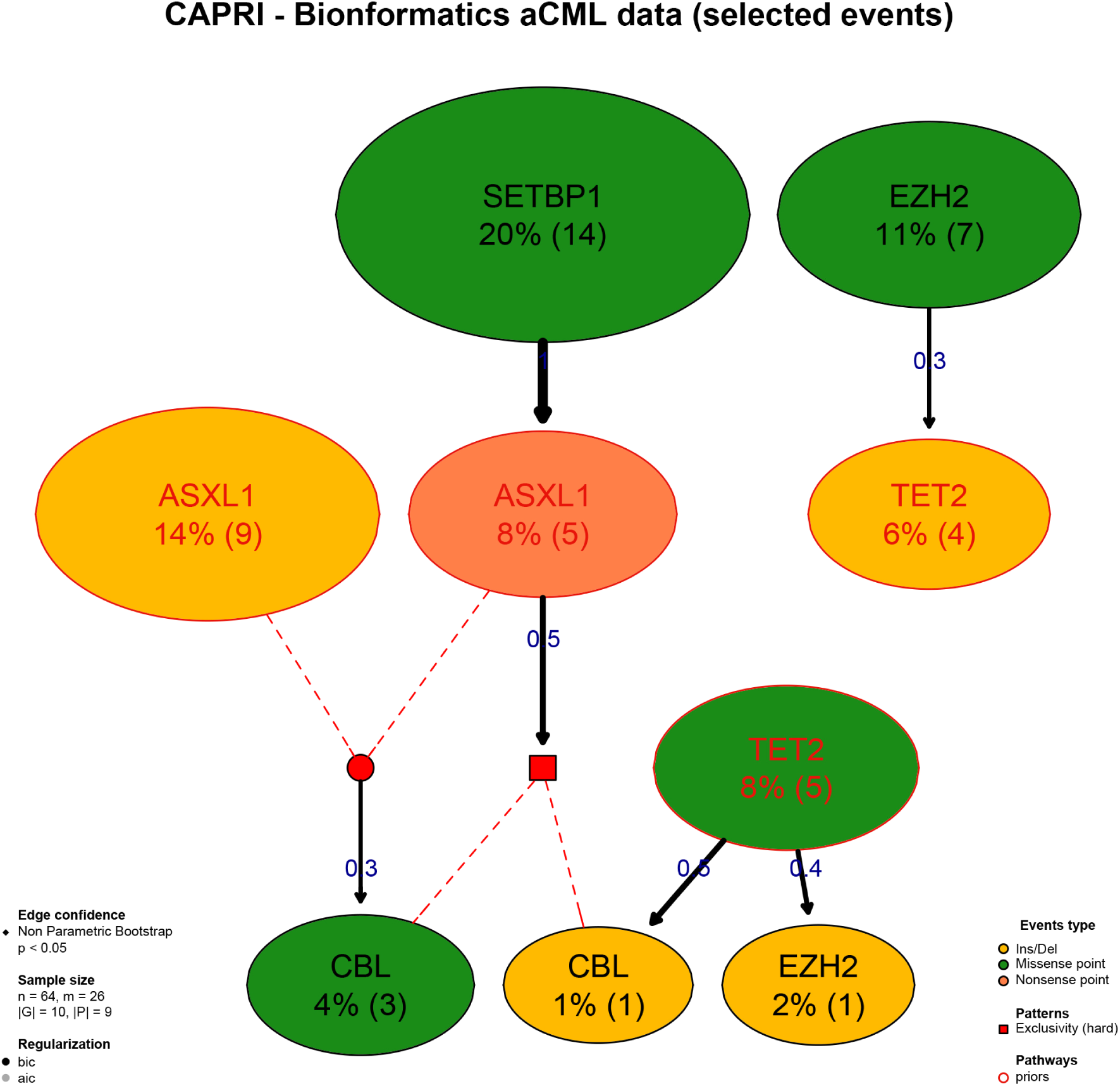
Reconstruction by capri and Bootstrap. Result of the reconstruction by capri on the input dataset with the assessment by non-parametric bootstrap.

~~~
> model.boot = tronco.bootstrap(model, nboot = 10)
Executing now the bootstrap procedure, this may take a long time…
Expected completion in approx. 00h:00m:03s
*** Using 7 cores via “parallel”
*** Reducing results
~~~

`Performed non-parametric bootstrap with 10 resampling and 0.05 as pvalue for the statistical tests.`

~~~
> tronco.plot(model.boot,
                 fontsize = 13,
                 scale.nodes = 0.6,
                 regularization = “bic”,
                 confidence = c(‘npb’),
                 height.logic = 0.25,
                 legend.cex = 0.5,
                 pathways = list(priors = gene.hypotheses),
                 label.edge.size = 10)
*** Expanding hypotheses syntax as graph nodes:
*** Rendering graphics
Nodes with no incoming/outgoing edges will not be displayed.
Annotating nodes with pathway information.
Annotating pathways with RColorBrewer color palette Set1.
Adding confidence information: npb
RGraphviz object prepared.
Plotting graph and adding legends.
~~~

We now conclude this analysis with an example of inference with the caprese algorithm. As caprese does not consider any pattern as input, we use the dataset shown in Figure 3. These results are shown in Figure 9.

**Figure 9:**
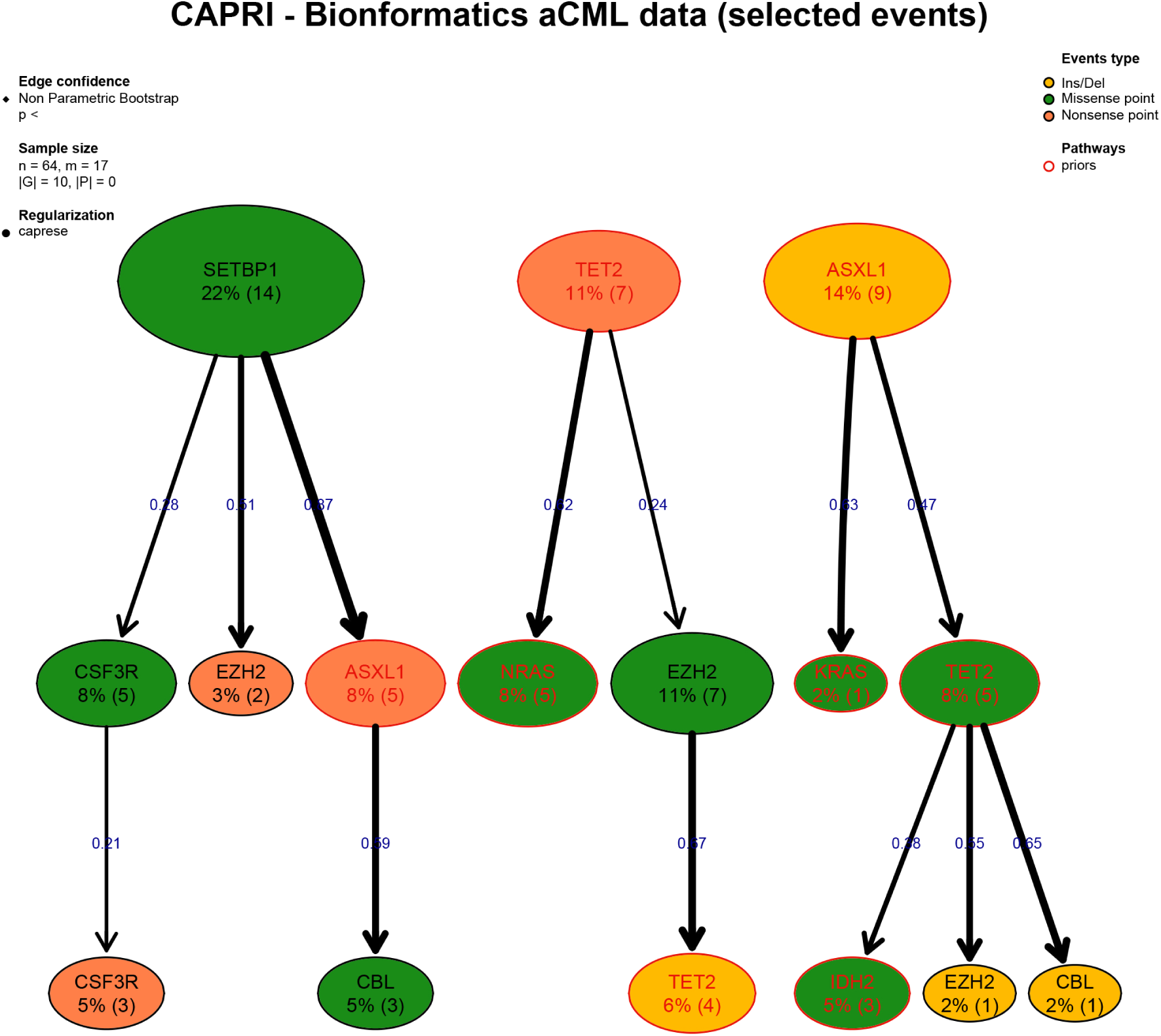
Reconstruction by caprese and Bootstrap. Result of the reconstruction by caprese on the input dataset with the assessment by non-parametric bootstrap.

~~~
> model.boot.caprese = tronco.bootstrap(tronco.caprese(hypo))
*** Checking input events.
*** Inferring a progression model with the following settings.
Dataset size: n = 64, m = 17.
Algorithm: CAPRESE with shrinkage coefficient: 0.5.
The reconstruction has been successfully completed in 00h:00m:00s
Executing now the bootstrap procedure, this may take a long time…
Expected completion in approx. 00h:00m:00s
~~~

`Performed non-parametric bootstrap with 100 resampling and 0.5 as shrinkage parameter.`

~~~
> tronco.plot(model.boot.caprese,
                 fontsize = 13,
                 scale.nodes = 0.6,
                 confidence = c(‘npb’),
                 height.logic = 0.25,
                 legend.cex = 0.5,
                 pathways = list(priors = gene.hypotheses),
                 label.edge.size = 10,
                 legend.pos = “top”)
*** Expanding hypotheses syntax as graph nodes:
*** Rendering graphics
Nodes with no incoming/outgoing edges will not be displayed.
Annotating nodes with pathway information.
Annotating pathways with RColorBrewer color palette Set1.
Adding confidence information: npb
RGraphviz object prepared.
Plotting graph and adding legends.
~~~

## 5 Conclusions

We have described tronco, an R package that provides sequels of state-of-the-art techniques to support the user during the analysis of cross-sectional genomic data with the aim of understanding cancer evolution. In the current version, tronco implements caprese and capri algorithms for cancer progression inference together with functionalities to load input cross-sectional data, set up the execution of the algorithms, assess the statistical confidence in the results and visualize the inferred models.

### Financial support

MA, GM, GC, AG, DR acknowledge Regione Lombardia (Italy) for the research projects RetroNet through the ASTIL Program [12-4-5148000-40]; U.A 053 and Network Enabled Drug Design project [ID14546A Rif SAL-7], Fondo Accordi Istituzionali 2009. BM acknowledges founding by the NSF grants CCF-0836649, CCF-0926166 and a NCI-PSOC grant.

## Footnotes

1 For capri the *n* actually depends on the structural complexity of the input “patterns”, i.e., of the boolean formulæ employed in the \lifting operation”; more information of this in [12].

